# Curation of causal interactions mediated by genes associated to autism accelerates the understanding of gene-phenotype relationships underlying neurodevelopmental disorders

**DOI:** 10.1101/2023.01.09.523265

**Authors:** Marta Iannuccelli, Alessandro Vitriolo, Luana Licata, Cristina Cheroni, Luisa Castagnoli, Giuseppe Testa, Gianni Cesareni, Livia Perfetto

**Affiliations:** Department of Biology, University of Rome Tor Vergata, Via della Ricerca Scientifica, 00133, Rome, Italy; Neurogenomics, Human Technopole, Viale Rita Levi-Montalcini 1, 20157, Milan, Italy; Department of Experimental Oncology, European Institute of Oncology IRCCS, Via Adamello 16, 20139, Milan, Italy; Department of Oncology and Hemato-Oncology, University of Milan, Via Santa Sofia 9, 20122, Milan, Italy; Department of Biology and Biotechnology “C.Darwin”, Sapienza University of Rome, Rome, Italy

**Author notes:** Equal contribution (Marta Iannuccelli, Alessandro Vitriolo).

## Abstract

Autism spectrum disorder (ASD) comprises a large group of neurodevelopmental conditions featuring, over a wide range of severity and combinations, a core set of manifestations (restricted sociality, stereotyped behavior and language impairment) alongside various comorbidities. Common and rare variants in several hundreds of genes and regulatory regions have been implicated in the molecular pathogenesis of ASD along a range of causation evidence strength. Despite significant progress in elucidating the impact of few paradigmatic individual loci, such sheer complexity in the genetic architecture underlying ASD as a whole has hampered the identification of convergent actionable hubs hypothesized to relay between the vastness of risk alleles and the core phenotypes. In turn this has limited the development of strategies that can revert or ameliorate this condition, calling for a systems-level approach to probe the cross-talk of cooperating genes in terms of causal interaction networks in order to make convergences experimentally tractable and reveal their clinical actionability. As a first step in this direction, we have captured from the scientific literature information on the causal links between the genes whose variants have been associated with ASD and the whole human proteome. This information has been annotated in a computer readable format in the SIGNOR database and is made freely available in the resource website. To link this information to cell functions and phenotypes, we have developed graph algorithms that estimate the functional distance of any protein in the SIGNOR causal interactome to phenotypes and pathways. The main novelty of our approach resides in the possibility to explore the mechanistic links connecting the suggested gene-phenotype relations.

## INTRODUCTION

### Vulnerability loci in neurodevelopmental disorders (NDDs)

The identification of vulnerability loci that underlie neuropsychiatric disorders has made considerable progress over the past two decades (Bray & O’Donovan, 2018), ushering, on the one hand, in the realization of their “daunting polygenicity”, with a large number of susceptibility loci contributing to genetic backgrounds of variable and often shared vulnerability across neuropsychiatric categories (Brainstorm Consortium, 2018). On the other hand, this effort has contributed to the identification of high penetrance, often de novo, monogenic variants (both mutations and copy number variations). Together, these insights have been increasingly breaking down highly prevalent diagnostic categories into, respectively, gradients of risk loadings or a myriad of bona fide rare conditions. This trend is poised to increase, as evident from the recent estimate that more than 1,000 genes that are associated to different extents with neuro-developmental disorders have not been described yet (Kaplanis *et al*, 2020).

In the case of Autism Spectrum Disorder (ASD), different kinds of studies, from both human cohorts and model organisms, have been providing a massive knowledge basis on the genes associated with this condition (De Rubeis *et al*, 2014; Grove *et al*, 2019; Gilman *et al*, 2011; Satterstrom *et al*, 2020; Weiner *et al*, 2017; Fu *et al*, 2022; Cheroni *et al*, 2020). Results of GWAS studies and additional data have been combined by the curators of the Simons Foundation Autism Research Initiative (SFARI) database to list several hundred genes implicated in autism susceptibility. The SFARI resource (Banerjee-Basu & Packer, 2010)) associates to the listed genes a score, ranging from 1 (high confidence) to 3 (only suggestive evidence), reflecting the strength of the evidence linking them to ASD (https://gene.sfari.org/). In addition, the syndromic category (S) includes mutations that are associated with a substantial degree of increased risk but are not required for an ASD diagnosis.

Typically, large gene lists are analyzed using methods such as, for instance, overrepresentation analysis (ORA) or protein interaction network analysis. These approaches are useful to gain insight into the biological functions associated with the genes in the query list and provide hints about the cellular processes whose disruption may contribute to a phenotype of interest. Briefly, methods based on over-representation analysis rely on pathway annotation (Falcon & Gentleman, 2007; Kanehisa *et al*, 2017; Gillespie *et al*, 2022; Khatri *et al*, 2012) or ontology vocabularies such as the Gene Ontology (Ashburner *et al*, 2000) to investigate whether a gene list is significantly enriched in genes annotated to any given pathway or function. When applying such approaches to the SFARI gene list, a significant enrichment in genes annotated to synaptic regulation and chromatin remodeling is observed (Vitriolo *et al*, 2019).

Network representation of biological complexity and graph theory, on the other hand, are playing an increasingly important role in dealing with the intricacy of human physiology and pathology and in limiting the noise that is inherent in large datasets (Barabási *et al*, 2011). Network approaches represent protein relationships as graphs connecting physically interacting proteins and build on the observation that related proteins (e.g. true hits from screening experiments or gene products mutated in the same disease) are more connected in molecular interaction networks than random proteins (Ideker *et al*, 2002; Barabási *et al*, 2011).

ORA and network representations, however, have some limitations. On one hand, ORA suffers from the limited annotation coverage in the reference databases as about 40% of the human proteome is not annotated to any pathway by Reactome and KEGG (Gillespie *et al*, 2022; Kanehisa *et al*, 2017) and, as such, they do not contribute to adding information to this analysis. In addition, pathway annotation is biased by curator decisions on whether to assign a protein to a pathway.

On the other hand, networks based solely on evidence of physical interactions, despite having the strength of high proteome coverage (Orchard *et al*, 2014; Oughtred *et al*, 2021), cannot provide information about the effects triggered by environmental cues or by genetic perturbations.

Conversely, networks where the edges are associated with additional causal information such as a direction and a sign are more informative as they allow one to make hypotheses on the causal consequences of the disruption of a protein activity on the function of downstream effectors. In recent years, a number of resources (Licata *et al*, 2019; Csabai *et al*, 2022) have undertaken an effort to manually capture signaling information from published articles and to represent it in a machine-readable format. The causal information captured by the SIGnaling Network Open Resource (SIGNOR), albeit still incomplete, has the highest coverage of published causal information represented according to the activity flow model (Junker *et al*, 2012) (Figure 1A).

**Figure 1.**
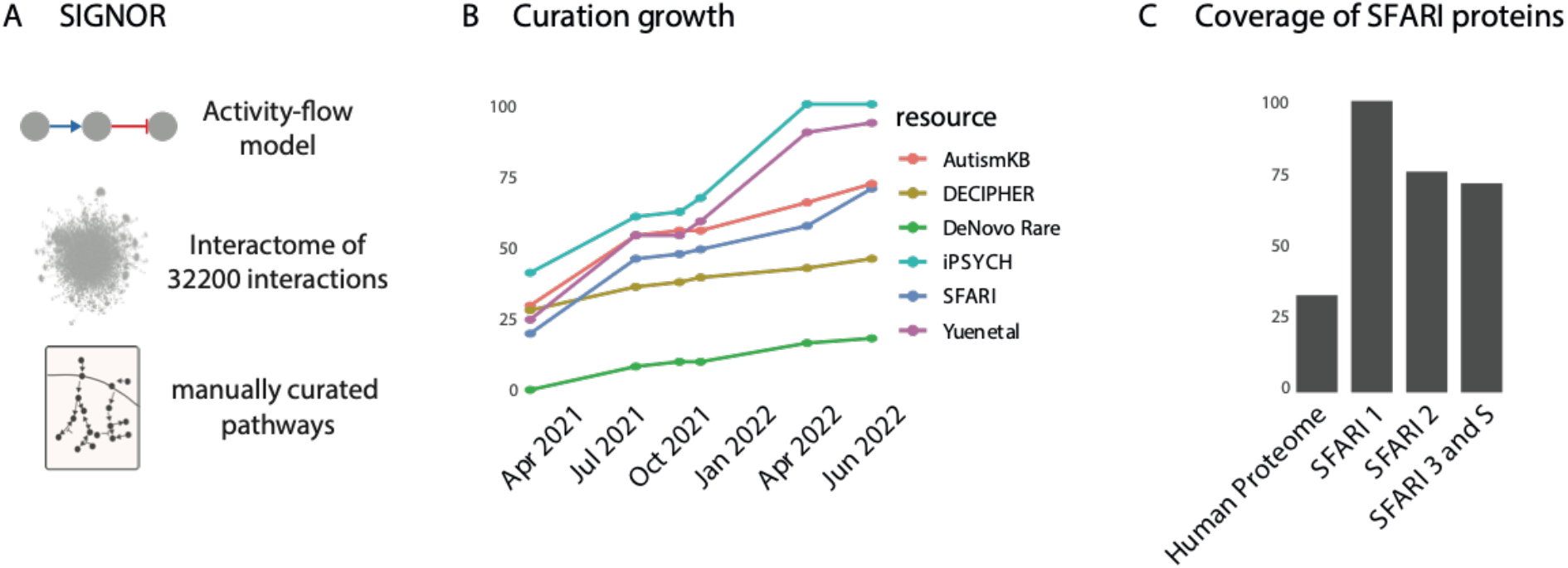
Curation of ASD-related causal relationships and pathways in the SIGNOR resource. **A)** Main features of the SIGNOR causal interaction database. Briefly, SIGNOR curators annotate interactions where the activity of one biological entity positively or negatively affects the activity of an interaction partner. The interactions are captured according to an ‘activity-flow’ model, where each interaction is binary, directed and signed (top panel). The interactions are integrated into a global cell network (middle panel). SIGNOR also annotates pathway maps that group set of interactions that modulate the activity of a biological process (bottom panel). B) Increase in annotation of genes associated to neuropsychiatric diseases by different resources: IPSYCH (Pedersen *et al*, 2018), AutismKB (Yang *et al*, 2018), DECIPHER (Firth *et al*, 2009) Yuen et al, (C Yuen *et al*, 2017), DeNovo rare (Guo *et al*, 2019). C) Coverage of SFARI gene lists in SIGNOR. SFARI 1,2,3 refer to lists of genes with different supporting evidence for association with Autism Spectrum Disorder as curated by the SFARI resource.

When in early 2021 this resource set out to provide a reference for causal interactions relevant for neuropsychiatric and neurodevelopmental diseases, only approximately 25% of the genes that had been associated with these disorders were part of the cell interactome in SIGNOR (Figure 1B).

We report here a curation effort, carried out over the past couple of years, aimed at increasing this coverage. In addition, to showcase the relevance of the curated dataset in dissecting the molecular mechanisms underlying neuropsychiatric diseases, we adapted our recently-developed computational strategy, here dubbed *ProxPath* (Perfetto *et al*, 2021; Iannuccelli *et al*, 2022). *ProxPath* exploits causal information annotated in SIGNOR to extend pathway and phenotype annotation in order to connect a larger fraction of autism-related proteins to a list of cellular pathways and phenotypes. This network-based approach contributes to identifying phenotypes that are ‘significantly close’ to a protein hit list.

## RESULTS

### Curation of causal interactions of genes and pathways associated to autism spectrum disorder

Curators of the SIGnaling Network Open Resource (SIGNOR) (Lo Surdo *et al*, 2022) manually annotate causal interactions according to an ‘activity-flow’ model (protein A up-/down-regulates protein B) (Cesareni *et al*, 2021) (Figure 1A). The resource captures signaling relationships between a variety of human biological entities, including biomolecules (proteins, macromolecular complexes, small molecules etc.), stimuli and phenotypes. Interactions in SIGNOR are assigned a significance score, ranging from 0.1 to 1 and form a large and intricate interactome of 9000 entities connected by 34,200 edges (November 2022) (Figure 1A). SIGNOR causal interactome is a large connected component with few satellite clusters (Lo Surdo *et al*, 2022). In parallel, SIGNOR curators also annotate pathways, which are subgraphs of the causal interactome providing a description of how a cell responds to specific environmental cues (Figure 1A). To date, SIGNOR annotates 114 manually curated pathways.

Here we set out to annotate causal interactions of prioritized ASD-related gene products and cellular pathways (Figure 1B-D). We took as a reference, for ASD-related genes, the dataset curated by the SFARI initiative (Banerjee-Basu & Packer, 2010). In February 2021 the SFARI gene resource listed 1003 ASD risk genes. At that time, 123/207 score 1 (High Confidence), 97/211 score 2 (Strong Candidate), 210/506 score 3 (Suggestive Evidence) and 41/79 score S (Syndromic) proteins were already included in the SIGNOR cell network. Since as much as 53% of the genes in the SFARI list were not annotated in SIGNOR, we initially compiled a ranked gene list based on SFARI gene score and prioritized for curation the genes with ascending score (from high to low confidence) that were also listed in other expert-curated resources(Pedersen *et al*, 2018; Yang *et al*, 2018; Firth *et al*, 2009; C Yuen *et al*, 2017; Guo *et al*, 2019).

By this approach, we were able to embed over 300 additional SFARI genes into the SIGNOR causal network and, as a result of this effort, 778 of the 1003 SFARI genes are now annotated in SIGNOR. Of these, the vast majority (770) are part of the large connected cell interaction network, whereas the remaining eight belong to small clusters that are not connected to the rest of the network (Supplementary Table 1). As shown in Figure 1C, after this curation effort, 99%, 77% and 71% of the SFARI score 1, 2 and 3 and S proteins, respectively, are now integrated into the cell causal interactome.

### ASD genes form a highly connected cluster in the causal network

In 2017, the group of Barabasi (Menche *et al*, 2017) provided evidence that patients affected by the same clinical conditions, despite being characterized by considerable genetic heterogeneity, show a high degree of homogeneity at the pathway level. This is consistent with the notion that the function of genes, found to be mutated in the same disease, often converge onto common signal transduction cascades (Pinto *et al*, 2014). We thus tested whether such pathway level convergence of ASD associated genes is observable in the SIGNOR causal network. Interestingly, 256 SFARI proteins form a large cluster fully connected by 363 directed causal edges (Figure 2). The *p-value* for such a cluster forming by chance was computed by counting the number of direct connections between SFARI genes in 1000 networks where the connections are randomized, maintaining node degree and edge direction (Iorio *et al*, 2016). The calculated *p-value* is in the order of 3*EXP-7 (Figure 2). Interestingly, overrepresentation analysis reveals that this cluster is enriched in proteins annotated with terms Axon Guidance, Circadian Clock, Dopaminergic Synapse, Glutamatergic Synapse and Neurotransmitters Release.

**Figure 2.**
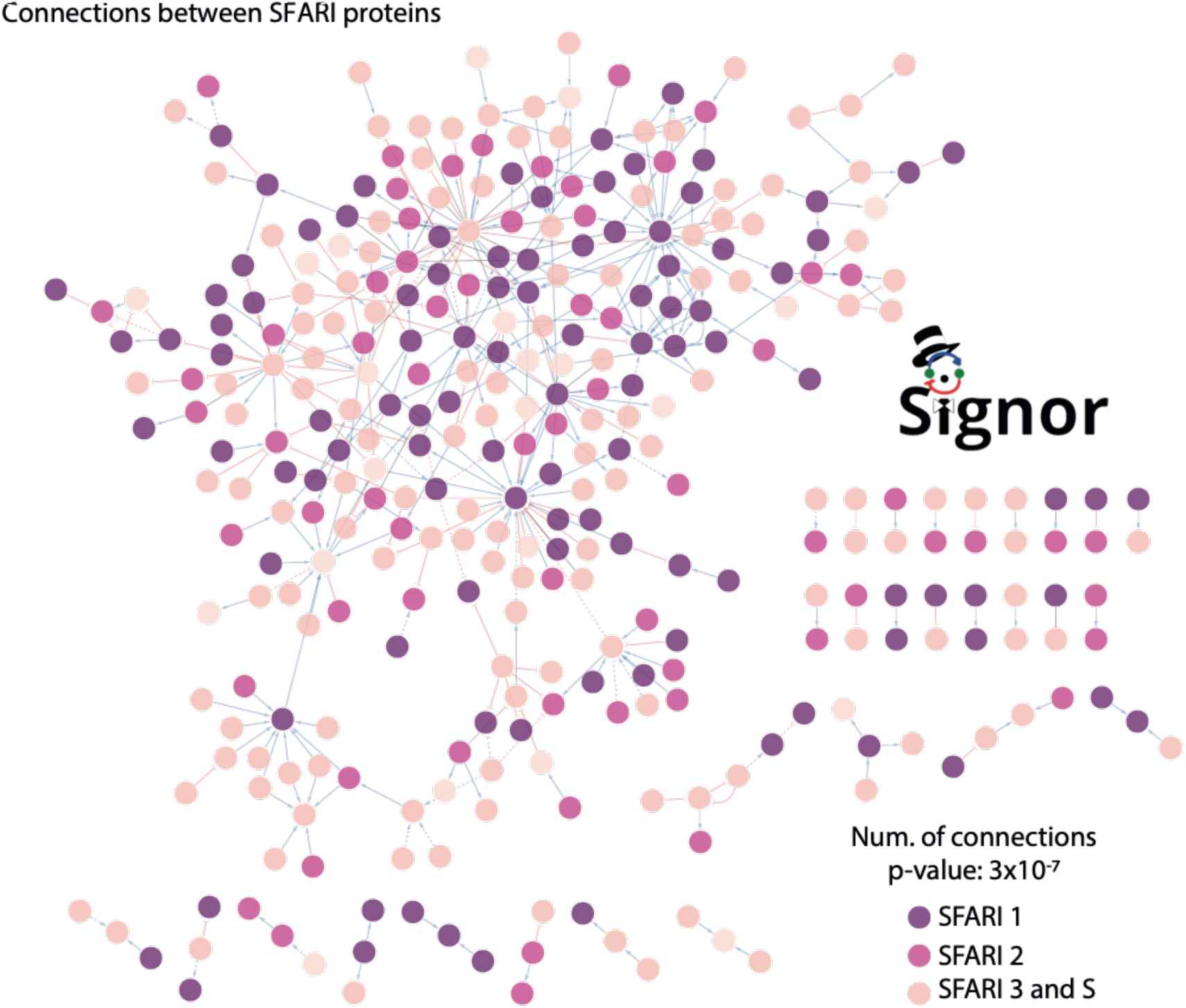
SFARI genes form a connected cluster in SIGNOR. Network of causal interactions directly linking SFARI proteins. The purple color grade of the nodes reflects the SFARI score classification as shown in the legend. Statistical significance of the clustering is measured as a p-value calculated by comparing the number of direct edges of the displayed network with those of 1000 randomly generated networks linking protein lists of identical size.

### Mapping ASD genes to pathways whose perturbation has been implicated in neurological disorders

In parallel to the gene and phenotype annotation work, we have also curated a list of 16 pathways that have been reported to be linked to ASD. They include signal transduction cascades governing neuron development and differentiation, synaptic assembly and transmission. In addition, we have also curated pathways that were found perturbed in ASD patients (e.g. Sex Hormone Biosynthesis (Janšáková *et al*, 2020) or WNT (Kwan *et al*, 2016)), or biological processes that emerged from the analysis of ASD-related genes (e.g. mRNA maturation) (Joo & Benavides, 2021). The curated pathways, with the exception of the ‘mRNA maturation’ pathway, form a single connected cluster (Figure 3).

**Figure 3.**
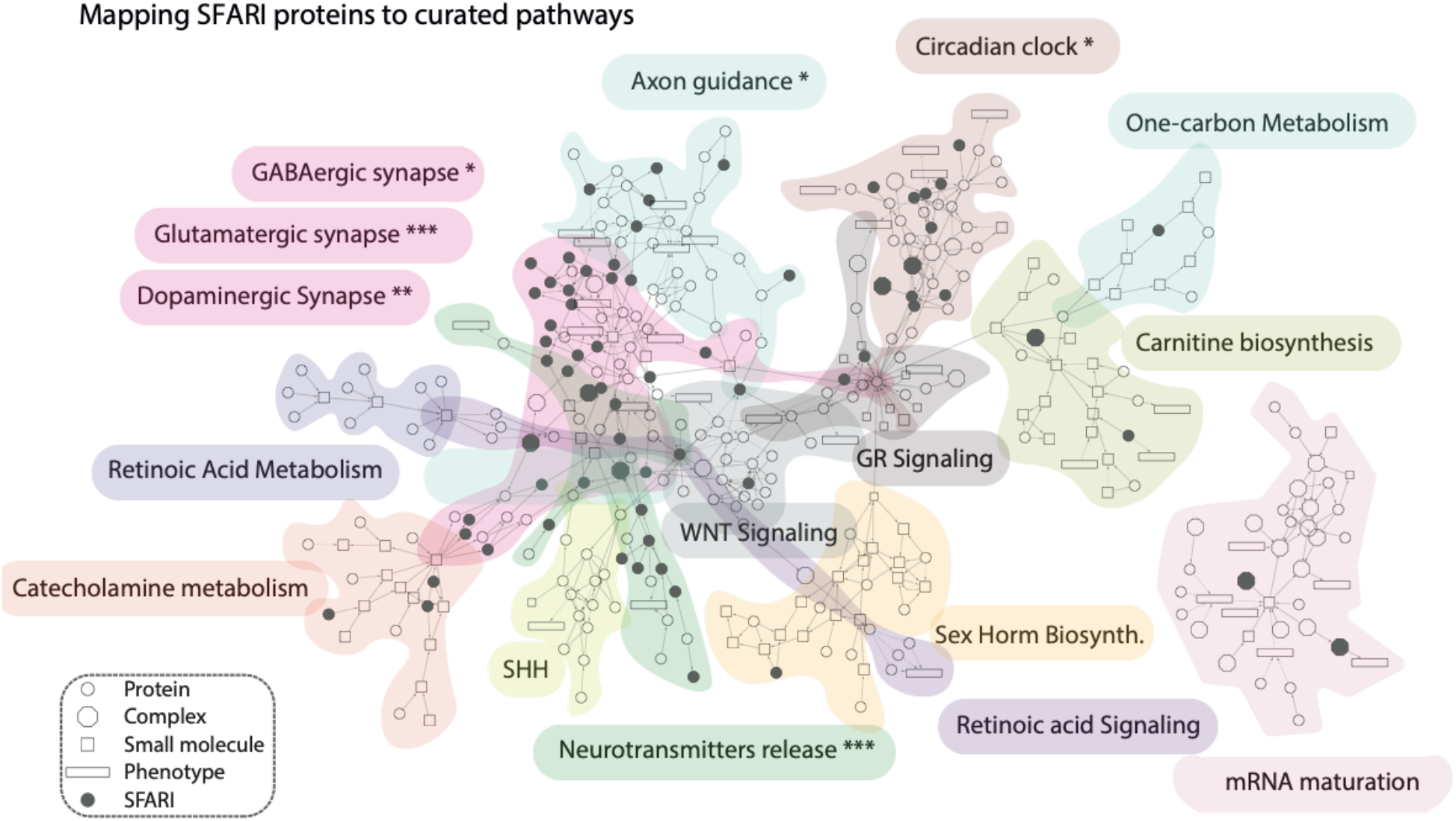
Mapping SFARI prioritized genes onto pathways of neurological relevance. The scaffold Network is obtained by merging pathway maps curated in SIGNOR. Nodes are biological entities: circles correspond to proteins, rectangles to phenotypes, squares to small molecules, octagons to protein complexes. Black nodes indicate SFARI proteins or complexes containing SFARI proteins. Significance of enrichment of SFARI genes in the considered pathways was evaluated by a Fisher’s exact test using as background all the proteins in SIGNOR (Bonferroni-corrected p-value: * <0.05, ** <0.01, *** <0.001). Mapping SFARI genes on such a graph (dark gray nodes) permits focusing on the pathways that are significantly enriched in the SFARI gene list.

As shown in Figure 3, SFARI proteins (black circles) preferentially participate (*p-value* < 0.05) in pathways involved in neurotransmitter release or synaptic transmission. Interestingly, the observation of a significant SFARI-protein enrichment in the circadian clock pathway provides support to the suggestion that dysregulation of the circadian rhythms plays a role in Autism Spectrum Disorder (Glickman, 2010). While this type of analysis is reminiscent of a conventional pathway enrichment analysis, it provides additional crucial information. The mapping of disease genes to pathways embedded in a causal network enables in fact the following: i) inspect pathway cross-talk, ii) formulate hypotheses on the mechanisms that are disrupted in the diseases and iii) provide suggestions on how to revert the disease phenotype by network intervention. The details of the network in Figure 4 can be inspected at https://www.ndexbio.org/viewer/networks/fbc6ec1f-fe96-11ec-ac45-0ac135e8bacf.

**Figure 4.**
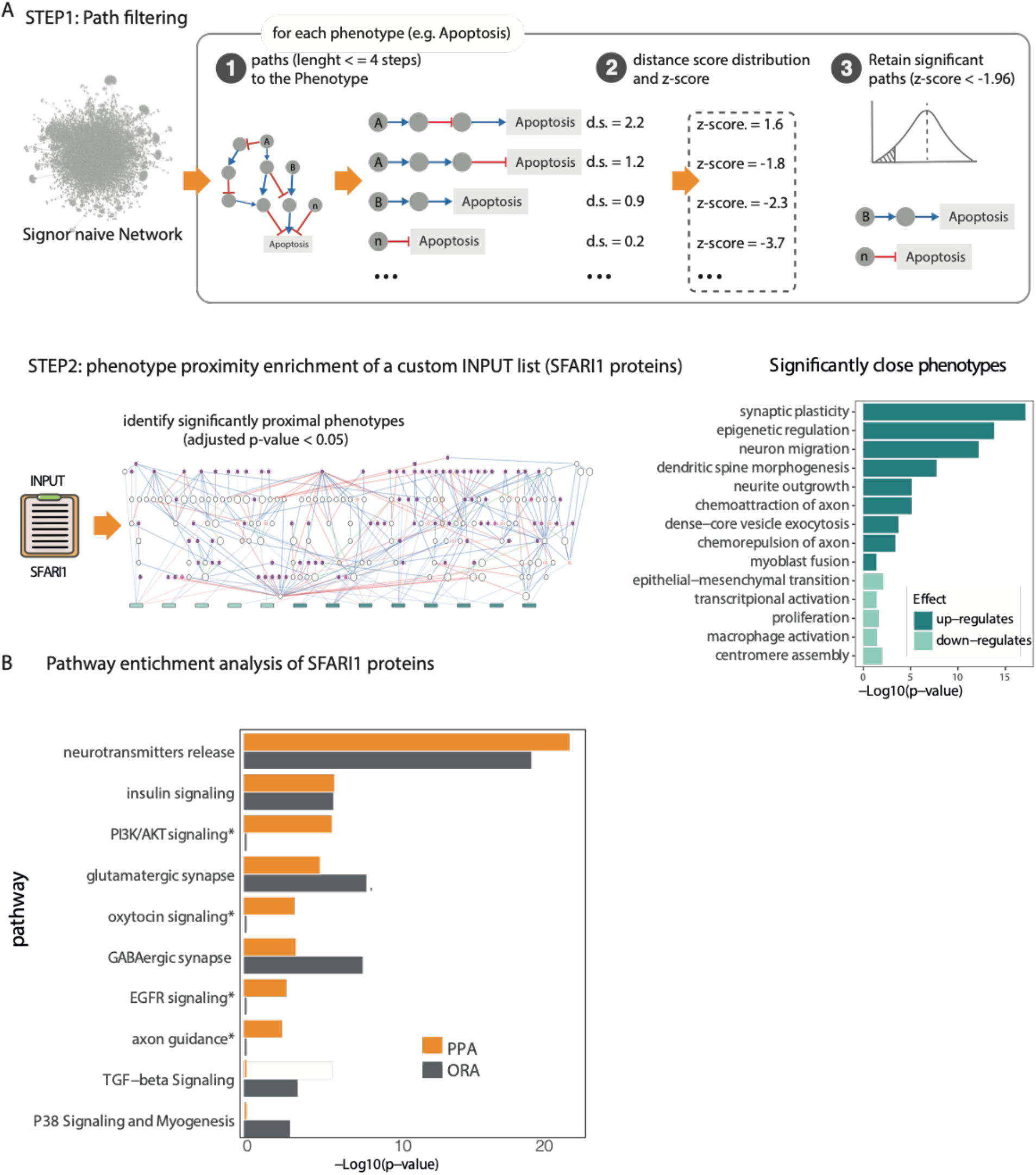
Estimating the regulatory impact of SFARI (score 1, SFARI1) proteins on phenotypes and pathways annotated in SIGNOR. (A) STEP1: The SIGNOR causal network is filtered to retain only significantly short paths impacting on phenotypes: STEP 1.1: for each protein-phenotype pair we calculate all the weighted distances in paths shorter than four steps; STEP1.2: for each phenotype we plot a distance distribution and for each path we calculate the Z-score; STEP1.3: Proteins with a Z-score lower than −1.96 (p-value <0.05) are considered to be likely to modulate phenotype expression. STEP2: significant paths extracted from STEP1 are used to determine which are the phenotypes that are significantly close to a custom list of proteins (e.g. SFARI1 proteins), via t-test statistical analysis. The results of the enrichment can be visualized as networks of causal interactions (left panel) or as bar diagram (right panel). (B) Comparison of enrichment analysis by over representation (ORA) and pathway proximity (PPA) considering as input the SFARI1 list of genes.

### Pathway and phenotype annotation extension

The global causal cell interactome curated in SIGNOR allows to connect each pair of biological entities via weighted and directed graph-paths (Figure 4). The causal network also links cell pathways and phenotypes to proteins that have the potential to modify their activities. Embedding pathways and phenotypes into the cell causal network allows one to walk along the directional edges of the network and to estimate a causal distance between a protein and a phenotype or a pathway. To support this type of analysis, we recently implemented *ProxPath*, an algorithm that, given a set of proteins, estimates its regulatory impact over phenotypes and pathways annotated in SIGNOR (Iannuccelli *et al*, 2022) (Figure 4B). *ProxPath* identifies short causal paths linking two graph nodes and estimates their *functional distance*. The approach considers the ‘trust score’ of each graph edge, thus allowing to define quantitatively if and how the activity of a protein has the potential to modulate the activity of a phenotype or a pathway. In the SIGNOR-network pathways are collections of causally connected nodes while phenotypes are individual nodes. Approximately 200 phenotypes and 114 pathways are presently embedded into the SIGNOR cell causal network.

We here describe two applications of *ProxPath:* the first measures the functional distance of an input gene list from individual target nodes (e.g. phenotypes), whereas the second computes the regulatory distance of an input list of genes to lists of target nodes (e.g. proteins that belong to a pathway).

The distribution of functional distances of proteins from a phenotype (or a pathway) depends on how central the phenotype or the pathway is in the causal graph. Thus, it is crucial to first normalize the potential impact of a certain protein on the modulation of the target entity. To this end, for each phenotype, *ProxPath* first plots a distribution curve of the weighted distances of all proteins from the phenotype (See also methods section). The activity of proteins with a distance-distribution Z-score smaller than −1.96 are considered as having a significant chance of impacting the phenotype or pathway (Figure 4A, step 1.3). This strategy allows us to extend the association of proteins to phenotypes and pathways in an unbiased manner and to make it quantitative and in that respect independent of curators’ decisions on the proteins to be associated to a pathway or a phenotype. By this approach we label each protein as significantly close to any of the SIGNOR phenotypes depending on whether the path distance has a Z-score <-1.96 in the distance distribution curve (Figure 4A). A similar approach is used to identify proteins that are significantly close to a pathway, as already described (Perfetto *et al*, 2021).

In essence, this approach allows to extend node annotation to nodes that are functionally close and relieves the approach from biases caused by curator decisions, thus enhancing the power of functional enrichment analysis. In support of this approach we compared the results of pathway enrichment of curator annotated pathways with that obtained after pathway expansion by causal pathway proximity (PPA), obtained by applying *ProxPath*. To this end, as outlined in the Methods section, we used the same input (the SFARI 1 gene list), background (the entire human proteome) and multiple-testing correction method. As shown in Figure 4B, both approaches identified ‘neurotransmitter release’, ‘insulin signaling’, ‘glutamatergic synapse’ and ‘gabaergic synapse’ as significantly enriched pathway annotations. However, differences were also observed as pathway proximity analysis revealed in addition the ‘PI3K/AKT signaling’, ‘oxytocin signaling’, ‘EGFR signaling’ and ‘axon guidance’ pathways, whose perturbations have already been described in autism spectrum disorders (Chen *et al*, 2014; Russo, 2014; McFadden & Minshew, 2013). In summary, our PPA approach has recapitulated the results from standard ORA, while partially compensating for the incomplete coverage of resources ORA depends on (Figure 4B).

We have also asked whether the SFARI 1 gene list is significantly enriched for proteins that are functionally close to any of the 200 phenotypes annotated in SIGNOR. To this end we generated 1000 lists of random proteins and computed a p-value for a random list having a number of proteins, significantly close to each phenotype, which is equal or larger than that observed in the SFARI 1 list. Although this strategy has also a certain degree of arbitrariness, it provides an independent estimate of gene-phenotype association. We confirm that the SFARI 1 gene list is enriched for genes that have the potential of positively modulating phenotypes related to synapsis assembly and function (Figure 4A). Furthermore, the approach also revealed an enrichment of genes involved in ‘epigenetic regulation’ and ‘dense core vesicle exocytosis’, processes which were already associated with autism spectrum disorders (Grafodatskaya *et al*, 2010; Lund *et al*, 2021).

In summary, these analyses show that networks of causal interactions are useful to underpin cellular processes or pathways that are associated to a list of gene products and can partially alleviate the lack of coverage in pathway resources. In addition, as the approach identifies the causal interactions linking the query proteins to a phenotype, it makes it possible to draw a graph detailing the molecular steps by which the proteins in the list may impact the phenotypic expression (Figure 4A-B). The complete network of causal interactions delineating the paths impacting enriched phenotypes can be inspected at https://www.ndexbio.org/viewer/networks/b7d7e952-fe97-11ec-ac45-0ac135e8bacf.

### Integrating poorly characterized proteins into the cell causal network provides information on their functions

Not all the proteins in the human proteome are equally well-annotated. The ‘Illuminating the Druggable Genome (IDG)’ project has developed a web-based platform (Pharos) that aggregates functional information captured by over 60 resources (Nguyen *et al*, 2017). Pharos uses a knowledge-based classification system to rank proteins according to the degree to which they are studied, as evidenced by a variety of features, thus helping to identify less characterized proteins. 5932 understudied proteins whose functions have been poorly, or not at all, characterized are labeled as ‘understudied’ and form the Tdark proteome.

75 SFARI proteins are part of the Tdark proteome and, as a consequence, hardly any experimental evidence can support generation of hypotheses on the mechanisms underlying their contribution to ASD (Figure 5 A). However, 24 of the 75 SFARI proteins that are classified as Tdark according to Pharos are part of the SIGNOR causal interactome and can link to nodes in the network, including phenotypes. As examples, four causal edges or fewer can link NUDCD2 to *cerebral cortex development*, TANC2 to *dendritic spine morphogenesis* and IRF2BPL to *secretory granules organization*, three phenotypes that have already been implicated in ASD (Lo & Lai, 2020; Padmakumar *et al*, 2019; Walsh *et al*, 2008) (Fig. 5B). These observations point to the potential of a strategy based on linking poorly characterized genes to the cell causal interactome to shed light on their function.

**Figure 5.**
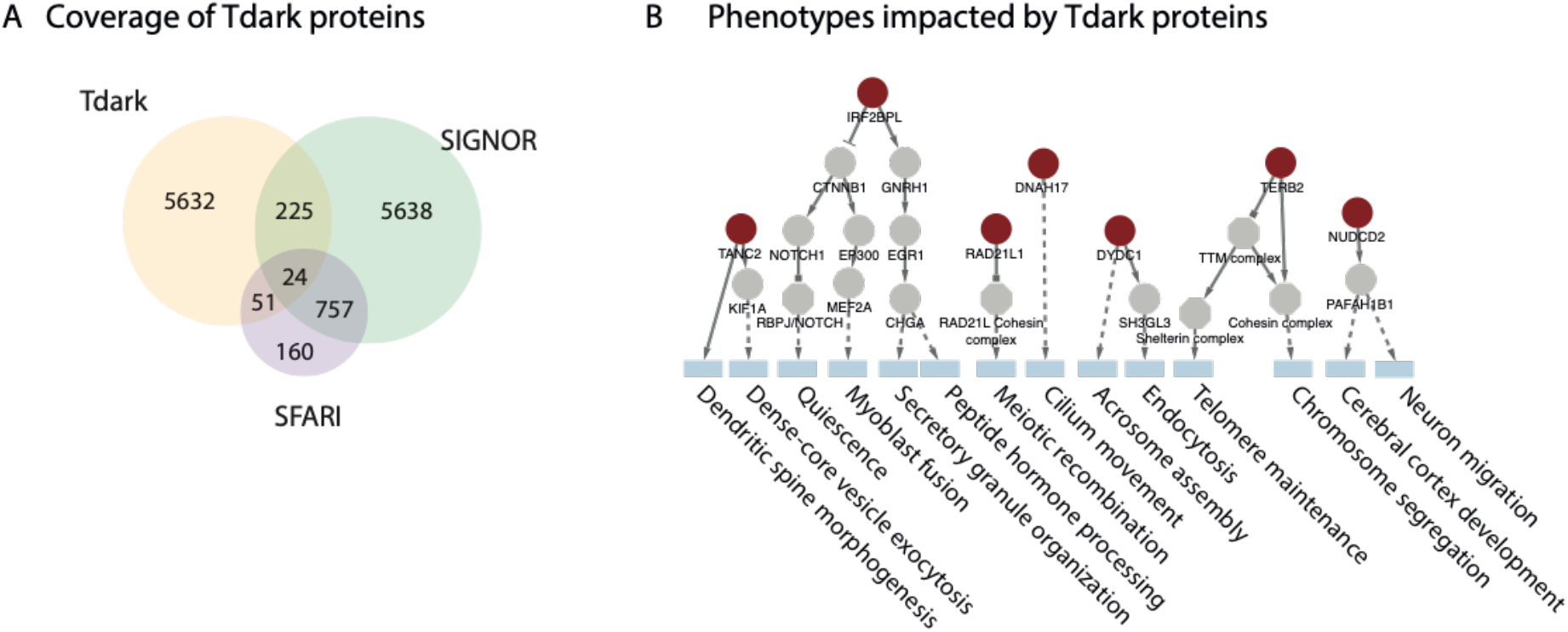
A) Overlap of proteins annotated in three resources: SIGNOR, SFARI and Tdark (Nguyen *et al*, 2017). B) Causal paths linking SFARI proteins to phenotypes annotated in SIGNOR. Tdark proteins are in red

### Adding support to rank genes with suggestive evidence of association to ASD

Over recent years, the SFARI-gene resource has collected genetic evidence from genomewide association studies to link human genes to autism spectrum disorders (ASD). Close to 1000 genes have been potentially linked to ASD. Thus, it is important to rank them according to the strength of the supporting evidence to prioritize candidate genes for time consuming follow up experiments. SFARI curators have grouped ASD genes into four score categories: 1 to 3, with decreasing levels of supporting evidence, and S (Syndromic genes).

As the large number of genes may cause the inclusion of false positives, especially in the categories with lower experimental support, additional and orthogonal scoring strategies may be useful to further prioritize genes. This is particularly true for those genes that are only included in the lists because of suggestive evidence.

We argue that genes whose activities underlie a given phenotype or whose disruption may contribute to a disorder are likely to be closely connected in a causal network. Thus, we took the list of high confidence SFARI genes (SFARI1) as a proxy of bona fide genes modulating the ASD phenotype. Next, for each gene in the proteome we used the distance from the closest gene in the SFARI 1 gene list as a proxy of their potential to be an ASD susceptibility gene. This approach generates a ranked gene list where the genes placed at the top of the list are the ones that receive more support from our network approach to be functionally connected to ASD (Supplementary Table 2).

### Common etiology of neuropsychiatric disorders

We next asked whether the value of our curation effort aimed at the integration of ASD genes into a cell causal network is not limited to the interpretation of the SFARI dataset and could more generally be valuable in neuropsychiatric studies. To this aim we monitored the annotation coverage of independently-defined lists of genes implicated in neuropsychiatric disorders, as reported by the Psychiatric Cell Map Initiative (Willsey *et al*, 2018) (Figure 6). The overall goal of this initiative is that of connecting genomic data to functional data (e.g. physical and genetic interactions) and ultimately to the clinic. Here we focus on autism spectrum disorders (ASD), intellectual disability (ID), epilepsy (EP), epileptic encephalopathies (EE) and Schizophrenia (SCZ). To avoid confusion the list of genes associated with ASD by the Psychiatric Cell Map Initiative will be indicated as ‘PCMI-ASD’.

**Figure 6.**
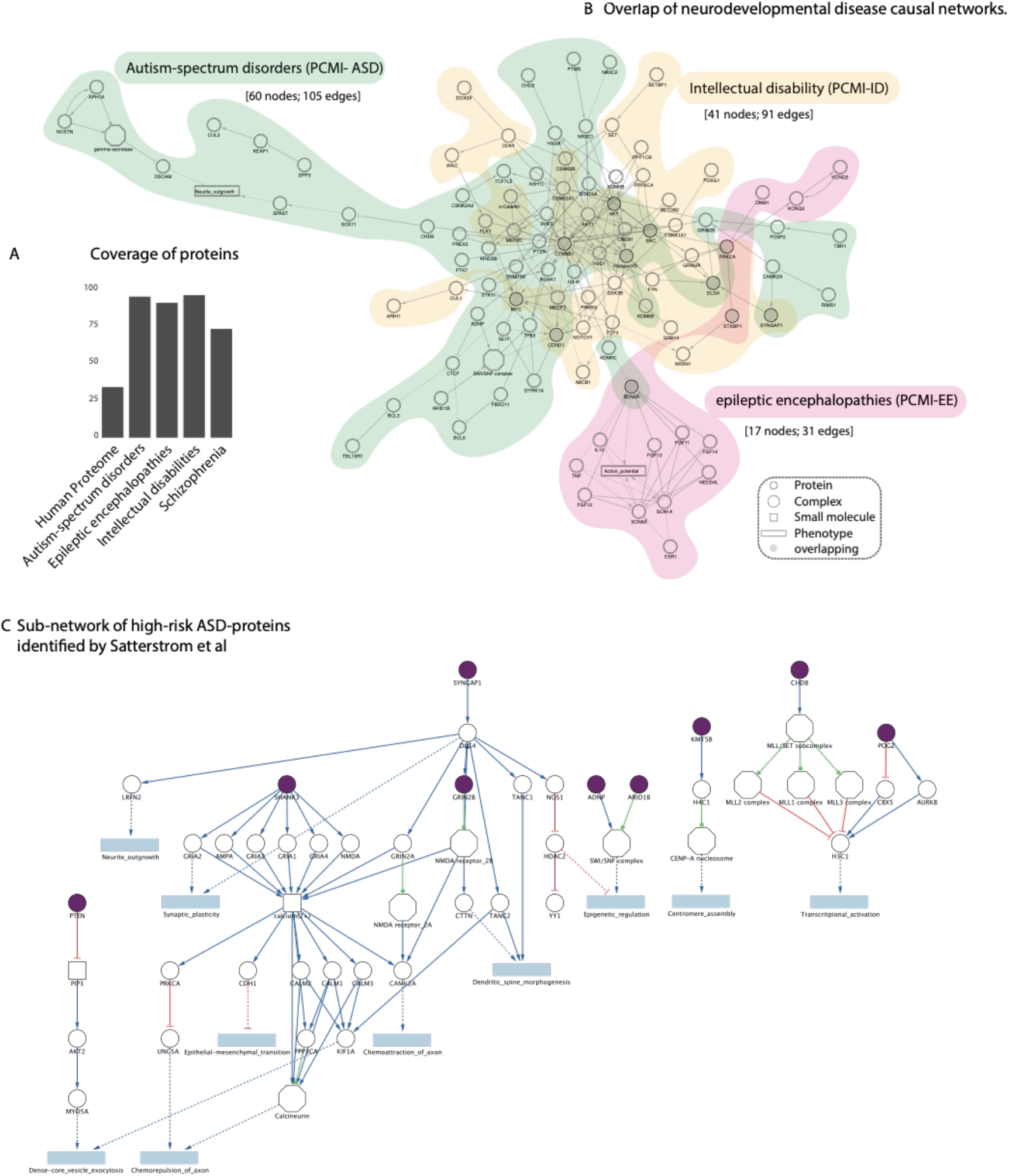
Overlap of neurodevelopmental disease causal networks. **(A)** Percent coverage of different neuropsychiatric-disease gene lists in SIGNOR. Schizophrenia; autism-spectrum disorders; epileptic encephalopathies; intellectual disabilities refer to the disease gene lists compiled by the Psychiatric Cell Map Initiative (Willsey *et al*, 2018). **(B)** Three disease networks are obtained by querying the SIGNOR resource with the neurodevelopmental disorders proteins annotated by the Psychiatric Cell Map Initiative to Autism spectrum disorders (PCMI-ASD), Intellectual disability (PCMI-ID), Epileptic encephalopathies (PCMI-EE). The three networks are shaded with green, yellow and pink backgrounds respectively. Nodes are biological entities: circles correspond to proteins, rectangles to phenotypes, squares to small molecules, octagons to protein complexes. Proteins that are common to more than one disease map are represented as black-circled nodes. **(C)** Sub-network of causal interactions extracted from the interactome in Figure 4A, linking 9 out of 13 high-risk ASD genes (in purple) (Satterstrom *et al*, 2020) to deregulated phenotypes (in blue).

We observe that our curation project also resulted in a high annotation coverage of these independently-defined gene lists (Figure 6A). Only 23 out of 47 proteins from PCMI-EE, PCMI-SCZ and PCMI-ID lists are also in the SFARI lists. Nevertheless, their coverage in the SIGNOR interactome is over 80%. Based on this observation, we speculate that the causal interactions annotated in SIGNOR are not limited to the SFARI dataset and to ASD but have also value for a broader range of neurodevelopmental and neuropsychiatric disorders.

To show this point, for each disease, we used the list of genes associated to four neuropsychiatric disorders, as annotated by the PCMI, to query the SIGNOR resource using the ‘connect + add bridge proteins’ search method (De Marinis *et al*, 2021). This method uses the causal information annotated in SIGNOR to draw networks indirectly connecting query proteins with bridge proteins. The three resulting graphs were displayed in a single layout forming a connected network (Figure 6B), No interaction paths could be drawn to link the PCMI-SCZ proteins. The PCMI-ID and PCMI-ASD maps appear to be highly interconnected. Fifteen proteins are common to the disease maps of more than one disorder (grey nodes in Figure 3). These include proteins that take part in the WNT and the Focal Adhesion pathways, suggesting that the deregulation of these biological processes might be implicated in more than one neurodevelopmental disease. These observations support the notion that the three disorders have a partially common molecular etiology.

### Leveraging the SIGNOR causal interactome to identify molecular convergences of ASD forms

We finally asked whether a causal network approach could recapitulate existing knowledge and provide hypotheses-generating observations. The genetic landscape of ASD is highly heterogeneous (Vitriolo *et al*, 2019), from highly penetrant variants (both mutations and copy number variations (CNV), often occurring *de novo*, to a large number of low single nucleotide polymorphisms (SNPs) interacting within individual genetic backgrounds (Grove *et al*, 2019)(Robinson *et al*, 2016) as well as environmental factors (Cheroni *et al*, 2020). This complexity translates into a Sorites paradox, where *monogenic* causes of ASD can be ascribed to rare *de novo* genetic variations of single genes (typically found with trio-exome sequencing experiments (Wright *et al*, 2018) that systematically occur among individuals with largely overlapping phenotypic traits (e.g. ADNP in Helsmortel Van der Aa Syndrome), while *polygenic* causes can be defined for the majority of cases, where no clear individual mutation is identified and the interaction of multiple low-risk SNPs, likely impinging on the same pathways touched by the monogenic cases, contributes to the onset of ASD. Large GWAS and exome-sequencing studies have highlighted the existence of genes that were found more frequently mutated in a cohort of more than 12K patients-derived samples (Satterstrom *et al*, 2020). This suggests that these genes play a preeminent role in shaping the condition phenotype. Moreover, thirteen of such genes fall within loci that are more recurrently hit by copy number variations. These thirteen genes are annotated in SIGNOR. Hence, we extracted from the network described in Figure 4A the subgraphs connecting these genes to significantly close phenotypes via causal relationships. Only 9 of the 13 genes could be connected to phenotypes via causal paths that are significantly short and are represented in the graphs in Figure 6C. By this approach we could identify two gene groups that recapitulate previous knowledge, a first one impinging on synaptic and neuronal activity and a second bearing on epigenetic regulation and transcriptional control. Although the two functional gene clusters are separate, they crosstalk as the postsynaptic scaffolding protein DLG4, which plays a critical role in synaptogenesis and synaptic plasticity, also impacts epigenetic regulation by negatively modulating the activity of HDAC2 via nitrosylation by nNOS. A second level of crosstalk is mediated by the chromatin regulators ADNP and POGZ that cooperate in modulating the expression of multiple clusters of synaptic genes, and whose mutation leads to a significant decrease of postsynaptic protein expression and glutamatergic transmission (Conrow-Graham *et al*, 2022; Markenscoff-Papadimitriou *et al*, 2021).

These cross-talks between the ‘synaptic’ and ‘epigenetic’ axes, as revealed by network analysis of the causal connections between disease genes and phenotypes, could explain the phenotypic similarity of neurodevelopmental disorders caused by germline mutations of regulators of these two axes (Gabriele *et al*, 2018). The synthetic perturbation of the activity of such regulators (Kampmann, 2020) could thus translate into similar electrophysiological endophenotypes, because of their functional proximity. However, given the centrality of chromatin remodeling into defining differentiation trajectories, we cannot exclude that this functional proximity could instead translate into differentiation biases and non-cell-autonomous effects, that globally result into similar phenotypes.

## DISCUSSION

Autism spectrum disorder (ASD) is a neurodevelopmental condition, frequently caused by mutations of synaptic and chromatin regulators, characterized by an early onset, and resulting in individually variable socio-cognitive impairments. The last decade has witnessed a major shift in our view of NDD as conditions potentially amenable to pharmacological interventions specifically geared to their causative mechanisms, as exemplified in the paradigm cases of Fragile X syndrome (Henderson *et al*, 2012), Down syndrome (Deidda *et al*, 2015) and 7q microduplication syndromes (7Dup) (Mihailovich *et al*, 2022; Lopez-Tobon *et al*, 2022). However, the difficulty in translating Fragile x insights from preclinical models to the human setting has also highlighted how key knowledge gaps still hamper such translational pipelines, a cautionary tale that becomes all the more relevant if we are to pursue the dauting polygenicity of ASD into rational subsets stratified by convergent and actionable molecular alterations. Towards this long-term goal, here we aimed at providing a resource for the community to streamline the identification of causal interactions between ASD vulnerability genes and hence of the most likely hubs of convergent dysregulation to be prioritized for experimental validation and translational pipelines. Specifically, we report two advances that help to elucidate the molecular mechanisms underlying the involvement of gene variants in disease onset and development.

First, expert curation has screened the literature and captured experimental information on the consequences of disrupting disease gene functions on the activity of downstream genes. This information has been integrated into a large cell causal network representing how the activity of gene products crosstalk and impact pathway expressions and phenotype manifestations. Thanks to this curation effort, over 90% of autism associated gene products are now integrated into the cell interactome and causally connected to the remaining gene products, pathways and phenotypes. The results of this project are now publicly available and can be freely explored by using tools offered by the SIGNOR resource website or downloaded for local analysis, in compliance with the FAIR principles (Wilkinson *et al*, 2016). The cell interactome can be navigated by graph algorithms and the mechanistic steps leading to functional crosstalk of any gene pair can be explored by navigating the network. Although in this project the focus of the curation were genes implicated in ASD, the network that we have assembled also includes many genes involved in other neuropsychiatric conditions and its utility can therefore be extended to all of them.

Second, we have developed graph algorithms that allow us to measure the functional distance of any gene from any pathway or phenotype, by leveraging the features of SIGNOR graph, which is directed and signed, and whose edges are weighted according to estimated supporting evidence.

Given the phenotypic and genetic complexity of autism spectrum disorders and the number of genes that have been found to be associated to these conditions, the literature abounds of suggestions of association of ASD genes to perturbation of practically any cell function (Choi *et al*, 2016; Baranova *et al*, 2021)(Jiang *et al*, 2000). Nevertheless, our approach provides independent evidence of some of these connections and offers the unprecedented opportunity to contextualize their interrelations. Here, we have used the above-mentioned algorithms to show that autism-associated genes are significantly more proximal to pathways and phenotypes involved in functions that underlie brain development. Moreover, our analyses have revealed significant functional connections with a sizable specific portion of cellular pathways, implicated in transcriptional and epigenetic regulation. Here, we claim that our approach not only allows us to connect genes to functions via causal links but it also provides suggestions on the mechanistic steps underlying these connections.

We have demonstrated the value of embedding disease genes into a causal cell interactome in order to formulate hypotheses on the molecular mechanisms leading to phenotypic perturbations caused by gene variants. It needs to be pointed out, however, that the cell causal interactome that is presently covered by the SIGNOR dataset is still incomplete and includes only 33% of the human proteome. As a consequence, some relevant causal links may be still missing and this may somewhat alter the analyses of gene-pathway distance. Furthermore, the cell causal network that we have presented here is obtained by integrating evidence from experiments performed in a variety of cell types, tissues or model systems. Many of these causal relationships may not be relevant for the function of the cell type that is affected during brain development in autism patients. Approaches to exploit single cell RNAseq datasets to develop cell type specific interactomes have been proposed (Mohammadi *et al*, 2019) and applications of these strategies to the SIGNOR dataset may help to assemble more biologically and clinically relevant cell interactomes. Nevertheless, our approach, while recapitulating existing knowledge, has revealed gene-phenotype/pathway connections that suggest mechanistic steps underlying such connections.

## MATERIALS AND METHODS

### Curation

Autism associated genes as annotated by the SFARI initiative were ranked according to their SFARI score and to the number of resources enumerating them for association to autism (Pedersen *et al*, 2018; Yang *et al*, 2018; Firth *et al*, 2009; C Yuen *et al*, 2017; Guo *et al*, 2019). Starting from highly ranked genes we have looked by standard Medline searches for published information reporting the consequences of disrupting gene function on downstream proteins. The curation of the reported experimental evidence was according to the SIGNOR database curation standards (Lo Surdo *et al*, 2022).

### Datasets

We accessed SIGNOR data using available RESTAPIs in June 2022 (De Marinis *et al*, 2021). At the time of writing SIGNOR annotates 32,200 interactions, between 8970 biological entities, 6700 of which are proteins. The SFARI genes list was downloaded from the SFARI resource in February 2021 (https://www.sfari.org/resource/sfari-gene/). Annotations as SFARI score 1 (207 proteins), 2 (211 proteins), 3 (506 proteins) and S (79 proteins) refer to the reliability of the annotation in the original resource, namely: category 1 (here indicated as SFARI score 1): High Confidence; category 2 (SFARI score 2): Strong Candidate; category 3 (SFARI score 3): Suggestive Evidence; category S (SFARI Syndromic protein). Syndromic mutations correlate with increased risk of ASD and are linked to additional characteristics not required for diagnosis.

Additional neurodevelopmental disorders protein lists were obtained from the Psychiatric Cell Map Initiative dataset (Willsey *et al*, 2018). 62 proteins are associated in these lists to autism spectrum disorders (PCMI-ASD), 28 to intellectual disability (PCMI-ID), 14 to epileptic encephalopathies (PCMI-EE) and 7 to schizophrenia (PCMI-SCZ).

### Network Analysis

Direct causal connections between SFARI genes were obtained by querying by the search mode ‘connect’ the SIGNOR database using the SIGNOR Cytoscape App(De Marinis *et al*, 2021). Statistical significance of the tendency to form clusters is measured as a p-value calculated by comparing the number of connections made by the SFARI proteins in 1000 randomly generated networks. Network randomization is obtained using the BiRewire method (Iorio *et al*, 2016) which preserves the functional characteristics of the nodes in the reference networks (i.e. their degrees and connection signs).

To obtain causal interaction graphs for neurodevelopmental diseases we used gene-disease association defined by the PCMI to query the SIGNOR database using the SIGNOR Cytoscape App (search mode: ‘connect + add bridge proteins’). We obtain the following networks: PCMI-ASD (60 nodes; 105 edges), PCMI-ID (41 nodes; 91 edges), PCMI-EE (17 nodes; 31 edges), PCMI-SCZ (0 nodes; 0 edges). We eventually merged the three networks using Cytoscape built-in function ‘merge’ to obtain an interactome of 103 nodes and 209 edges.

### ProxPath - estimating the Impact of a Protein on a phenotype

The *ProxPath* algorithm can estimate the regulatory impact of an input list of proteins, in our case SFARI (score 1) proteins, over phenotypes annotated in SIGNOR.

Significantly-proximal phenotypes are identified using a strategy adapted from Iannuccelli et al. (2022). The pipeline consists of two steps:

#### STEP1

Browsing the SIGNOR interactome to create Phenotype-proximity annotations. Briefly, we make use of the graph representation of the human causal network annotated in SIGNOR 3.0. For each phenotype, the algorithm retrieves all the directional paths (of length of four steps or fewer) connecting any protein in SIGNOR to the phenotype. As any step in the graph has a score (s), we define the distance between any two interacting proteins as d = 1-s and the path distance score (d.s.) of a path including more than two nodes as the sum of the distances of the edges forming the path. The lower the path distance score, the shorter is the ‘functional distance’ and more functionally relevant is estimated the path. Next, we classify as proteins having an influence on phenotype activity those proteins having Z-score in the path distance distribution lower than Z = −1.96 (p-value <0.05) The Z-score is computed over the distribution of path distance scores considering the distance between every protein in SIGNOR to a given phenotype. Nodes that cannot connect with four steps or fewer to phenotypes were arbitrarily assigned the highest path distance score in the distribution.

The paths connecting the query protein to a phenotype are characterized by a distance and by a sign that specifies whether the protein is inferred to have a positive or a negative effect on phenotype activity. Proteins connected by paths formed by an odd number of inhibitory steps are defined as inhibitors, otherwise are considered as activators.

Identification of the paths linking a query gene to the phenotype was programmatically implemented using the ‘all simple_paths’ function of the NetworkX module of the Python language (Hagberg *et al*, 2008). The function returns every short path linking any two nodes in an oriented graph. We set a length cut-off of 4 as input parameter in order to explore only pathways with a length that is shorter or equal to the chosen threshold. R scripting was used to run python scripts and to analyze results.

#### STEP2

phenotype proximity enrichment of SFARI (score 1) proteins. To identify the proximal phenotypes impacted by the SFARI 1 proteins and the resulting effect on the phenotype, we used the strategy outlined in the previous section to link 207 SFARI proteins to the up- or down-regulation of 196 phenotypes annotated in SIGNOR. We used the twosided t-test to assess whether the proportion of paths starting from a SFARI 1 protein and leading to the up-/down-regulation of a phenotype is significantly greater than the mean extracted from a randomized dataset. Briefly, we generated lists of 207 proteins that were randomly chosen among the proteins annotated in SIGNOR. 207 corresponds to the number of SFARI 1 proteins. We generated 1000 of such random genes lists (The entire human proteome was considered as a background of the analysis) and, for each, we evaluated the fraction of short paths (<= 4 steps) starting from the input protein and impacting the up- or down-regulation of each phenotype displayed in SIGNOR. We eventually charted the distribution of these fractions, performed a t-test and selected phenotypes that display a Benjamini-Hochberg-corrected p-value < 0.05.

### ProxPath - estimating the Impact of a Protein on a pathway

The *ProxPath* algorithm can estimate the regulatory impact of a list of INPUT proteins, in our case SFARI (score 1) proteins, on pathways annotated in SIGNOR. Significantly-proximal pathways are identified using a strategy already described in Perfetto et al. (2021). Briefly, we define a global distance, or proximity score, between a query protein and a pathway, by considering the paths between any protein in SIGNOR and all the proteins in the pathway. To this end we followed a four-step strategy: 1) We searched the cell causal interactome for paths of four steps, or less, linking any of the query proteins and each protein in the pathway-list; 2) We select, for each query protein-pathway protein pair, the path with the shortest distance (lower distance score, d.s. - see *Estimating the Impact of a Protein on a phenotype paragraph*); 3) If a query protein is connected to more than one protein in the pathway, we use an analogy with a parallel resistor and define the proximity score as the reciprocal of the sum of the reciprocals of the distances (d.s.) of each path linking the query protein to proteins in the pathway; 4) we classify as proteins having an influence on the pathway activity those query proteins having Z-score in the log(proximity score) distribution lower than Z = −1.96 (p-value <0.05) The Z-score is computed over the distribution of proximity scores considering the distance between every protein in SIGNOR to a given pathway. To identify the proximal pathways impacted by the SFARI 1 proteins, we used the statistical tests already described in the Estimating the Impact of a Protein on a phenotype paragraph.

### Pathway Over-representation analysis (ORA)

To perform classical pathway over-representation analysis of SFARI1 proteins, we considered all the pathways in SIGNOR, and then we extracted all the pathway members using SIGNOR REST APIs. For each pathway, we next performed Fisher-exact test using as input the SFARI1 proteins and as background the entire human proteome. P-values were corrected for multiple testing using Benjamini-Hochberg method.

